## Supplemental Material for "3D molecular phenotyping of cleared human brain tissues with light-sheet fluorescence microscopy"

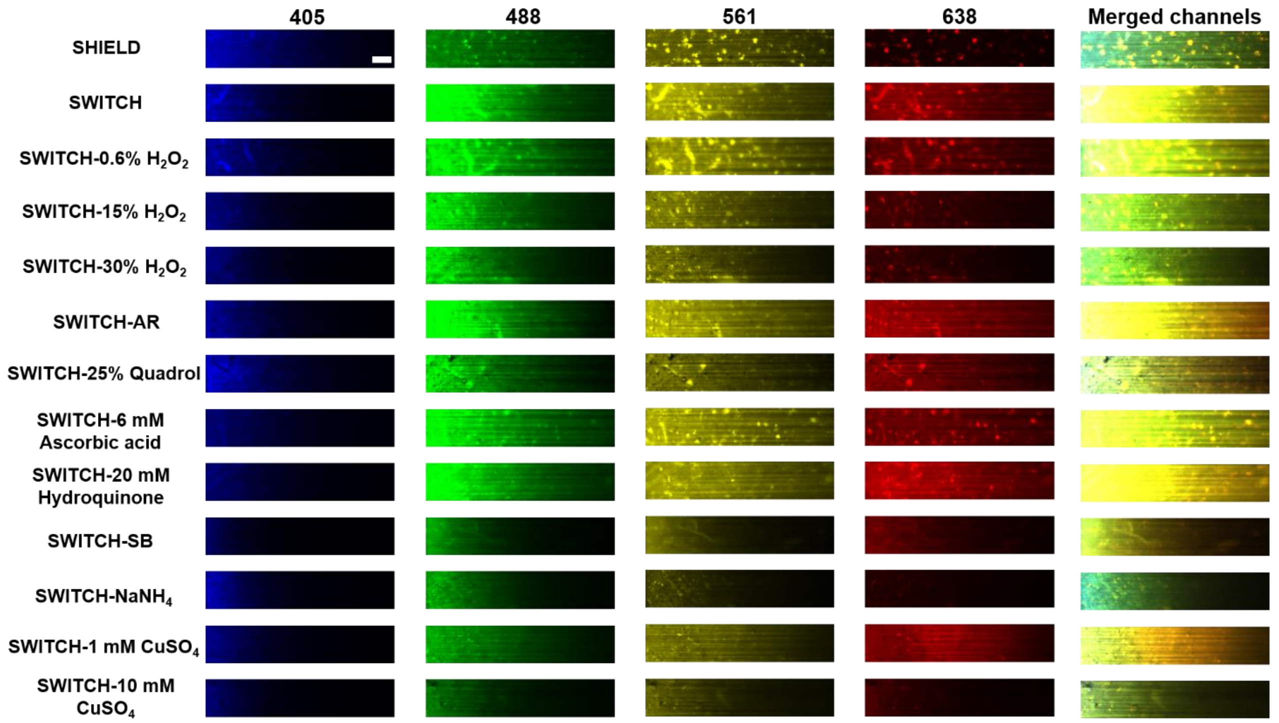


**Fig. S1. Characterization of SWITCH processed human brain slices treated with different autofluorescence elimination reagents.** Columns 1-4 correspond to the 4 different excitation lights; column 5 correspond to the merge channels for each treatment. The look-up tables (LUTs) are fixed for each treatment. Scale bar: 100 µm.

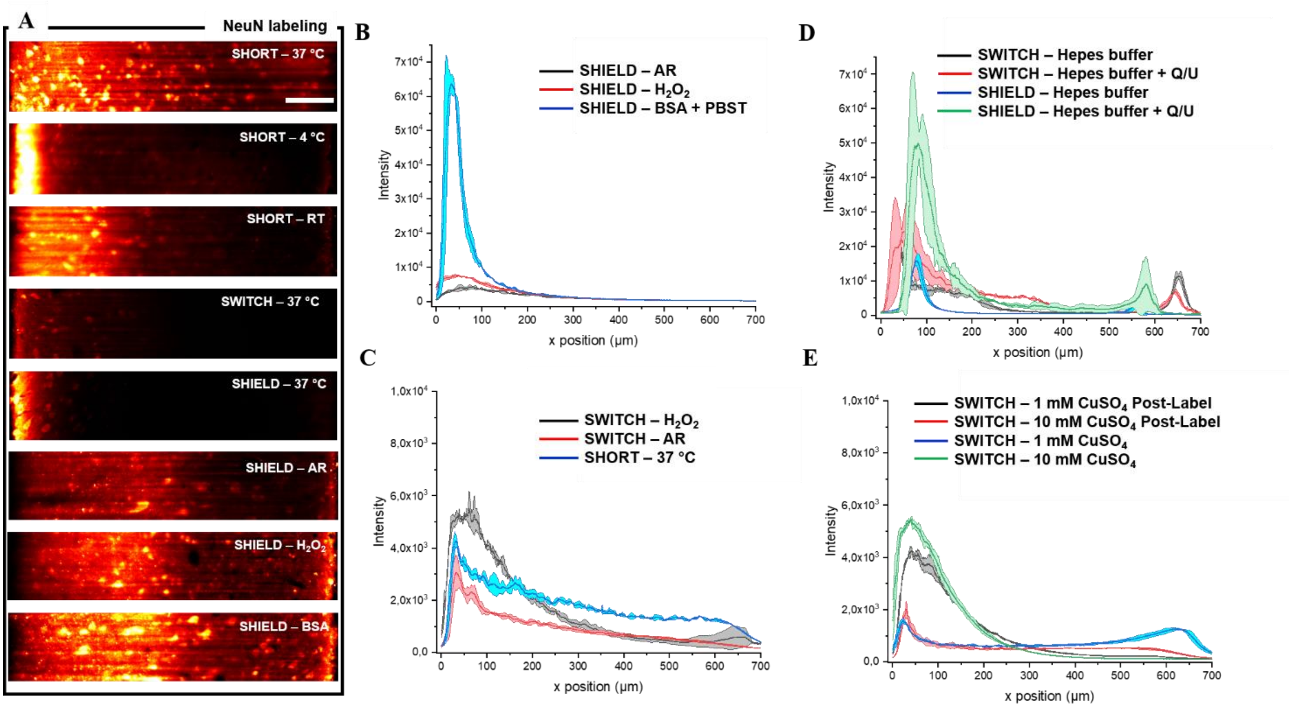


**Fig. S2. Effect of tissue processing (SWITCH, SHIELD, and SHORT), autofluorescence treatments ($CuSO_4$), and buffer on NeuN fluorescence labeling.** (**A**) High resolution imaging of NeuN labeling of different experiments shown in Fig. 1. Scale bar: 100 µm. (**B**) Profile plots of NeuN labeling of SHIELD-processed slices incubated with BSA + PBST, treated with AR or $H_2O_2$. (**C**) Comparison of SWITCH-processed slices treated with $H_2O_2$, AR or by combining both (SHORT). (**D**) Profile plots of NeuN antibody incubated with HEPEShepes buffer, or HEPES buffer supplemented with 2.5% Quadrol and 0.5 M urea in SWITCH/SHIELD-processed slices. (**E**) Profile plots of SWITCH-processed slices treated with 1 mM $CuSO_4$ after NeuN labeling (black and red profiles) or before NeuN labeling (blue and green profiles). Antibodies dilution for all experiments: NeuN 1:100; Alexa Fluor 647 1:500. Incubation temperature of 37 °C (N = 3). Excitation light 638 nm at laser power 5 mW.

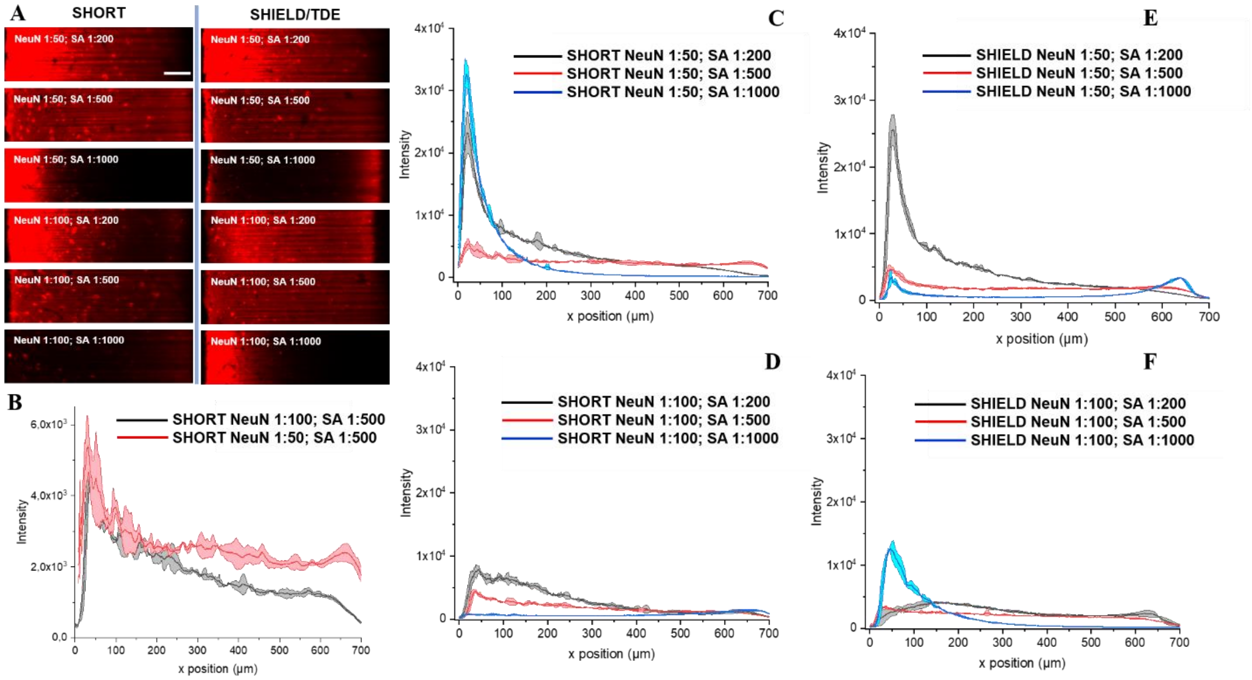


**Fig. S3. Antibody optimization of SHORT and SHIELD-processed slices.** (**A**) High resolution images acquired by LSFM of SHORT and SHIELD-processed slices, labeled for NeuN with Alexa Fluor 647 at different dilution (1:50 and 1:100 for NeuN; from 1:200 to 1:1000 for Alexa Fluor 647). Scale bar: 100 µm. (**B**) Profile plot along depth of the two best conditions of NeuN and Alexa Fluor 647 antibodies in SHORT-processed tissues. (**C** and **D**) Profile plots of different antibody dilutions of SHORT-processed slices. (**E** and **F**) Profile plot of different antibody dilutions of SHIELD-processed slices. Incubation temperature of 37 °C (n = 3). Excitation light 638 nm; laser power 5 mW.

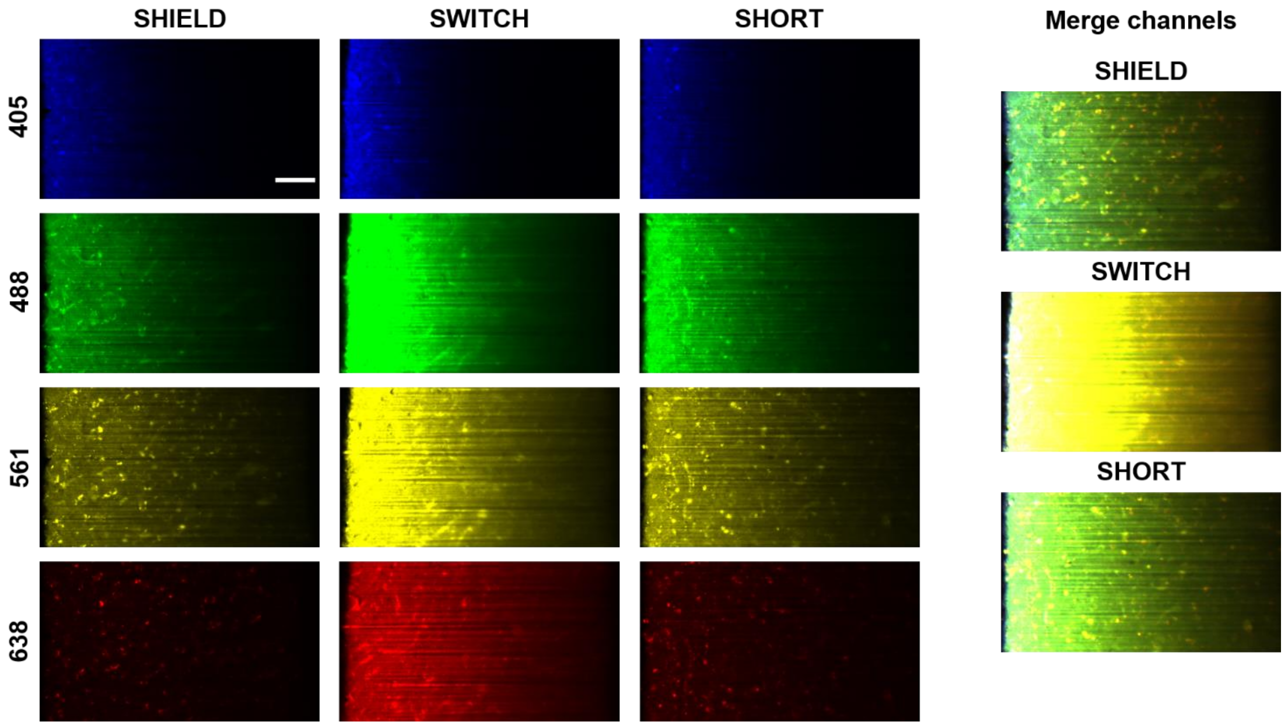


**Fig. S4. Comparison of autofluorescence signal between SHIELD, SWITCH and SHORT.** SWITCH-, SHORT-, and SHIELD-processed slices excited at 405, 488, 561 and 638 nm. SHIELD and SHORT show similar autofluorescence signal at 405, 561, and 638 nm, while SHORT shows higher intensity at 488 nm (see also Fig. 1A). Power: 5 mW. The LUTs are fixed for each treatment. Scale bar: 100 µm.

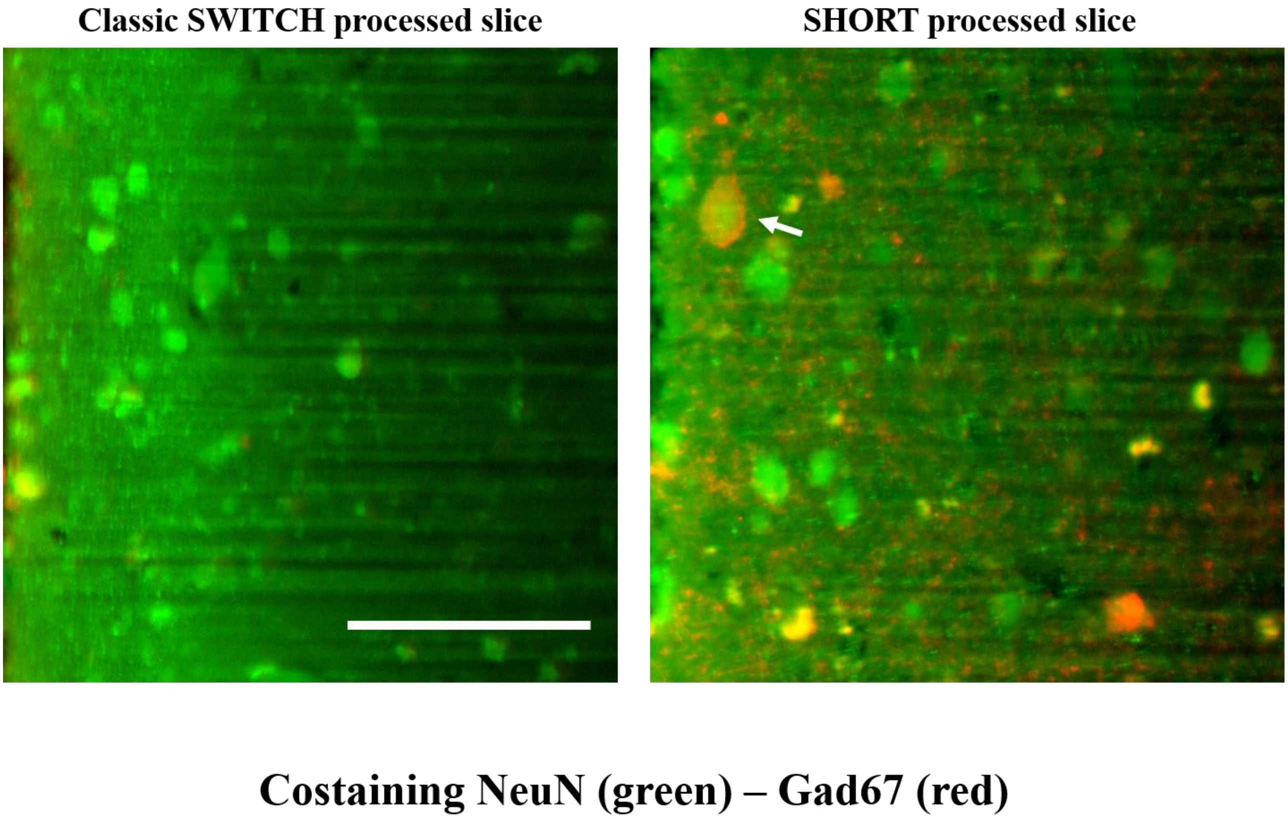


**Fig. S5. Comparison between the classic SWITCH protocol and SHORT for NeuN (green) - GAD67 (red) costaining.** The white arrow shows the high-resolution imaging of a typical costained neuron positive for NeuN and GAD67 in SHORT-processed slices. Scale bar: 100 µm. Excitation light: 488 nm and 638 nm.

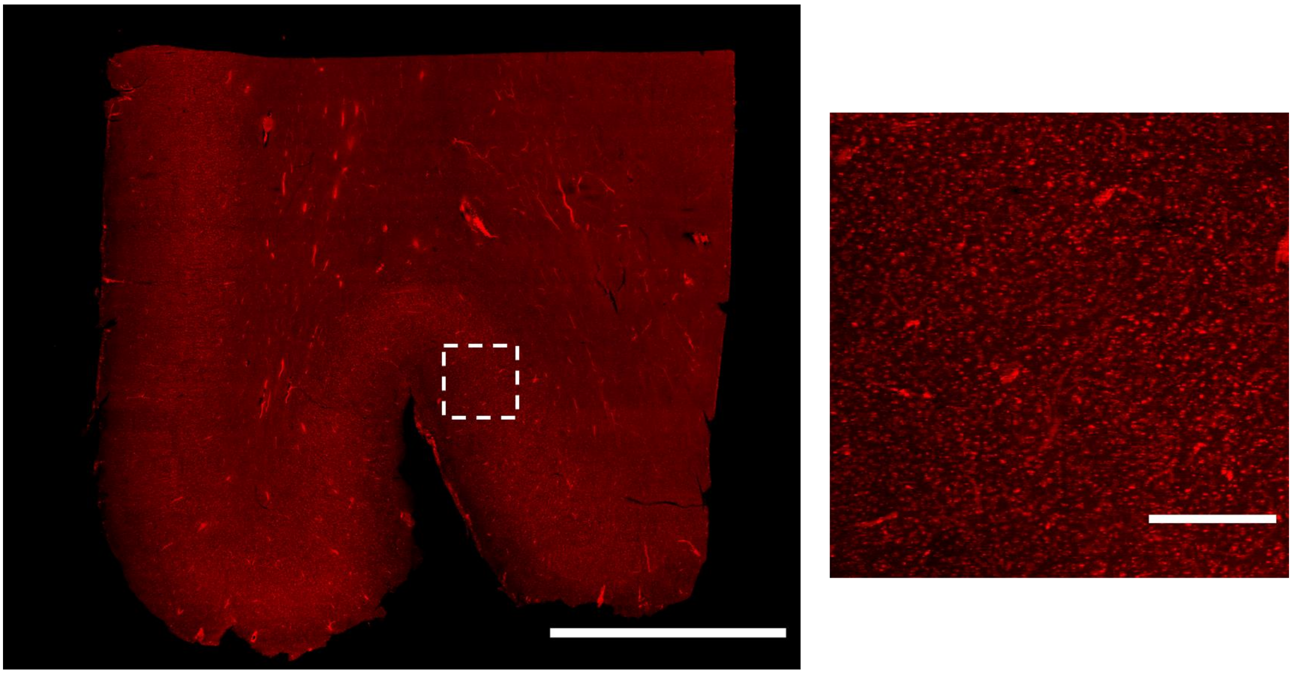

**Fig. S6. Propidium iodide (PI) allows an efficient nuclear labeling in SHORT-processed samples.** Human motor cortex labeled with PI and acquired using LSFM. Excitation light 561 nm. Scale bar: 5 mm. Magnified image of the nuclear staining, scale bar: 500 µm.

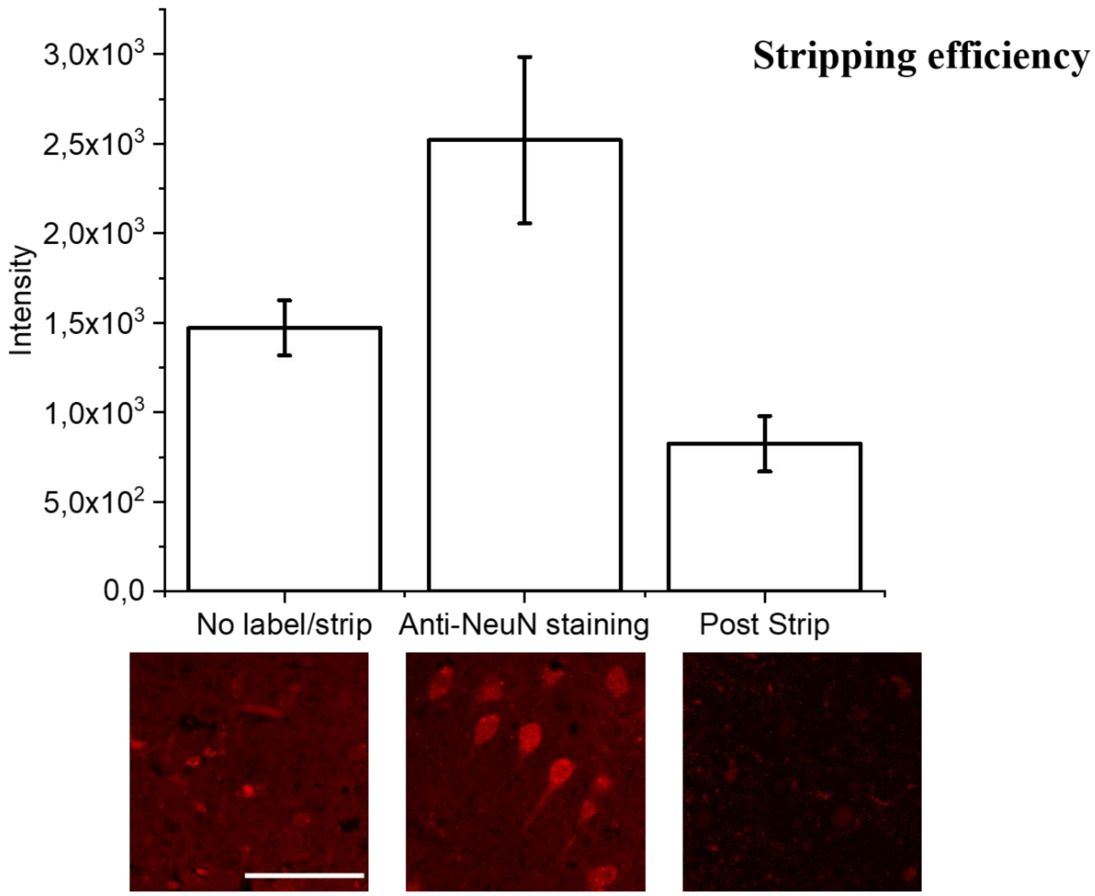


**Fig. S7. Stripping efficiency in SHORT-processed human slices.** Tissue sections were processed with SHORT and were acquired using a Nikon C2 laser-scanning confocal microscope. Then, the samples were immunostained for NeuN with AlexaFluor 568 and were re-acquired by the same microscope. The samples were then stripped using the elution buffer (see Methods) for 4 °h at 80 °C and were acquired again. Quantification of fluorescence intensity seen in n = 3 samples. Error bars: standard deviation. Scale bar: 50 µm.

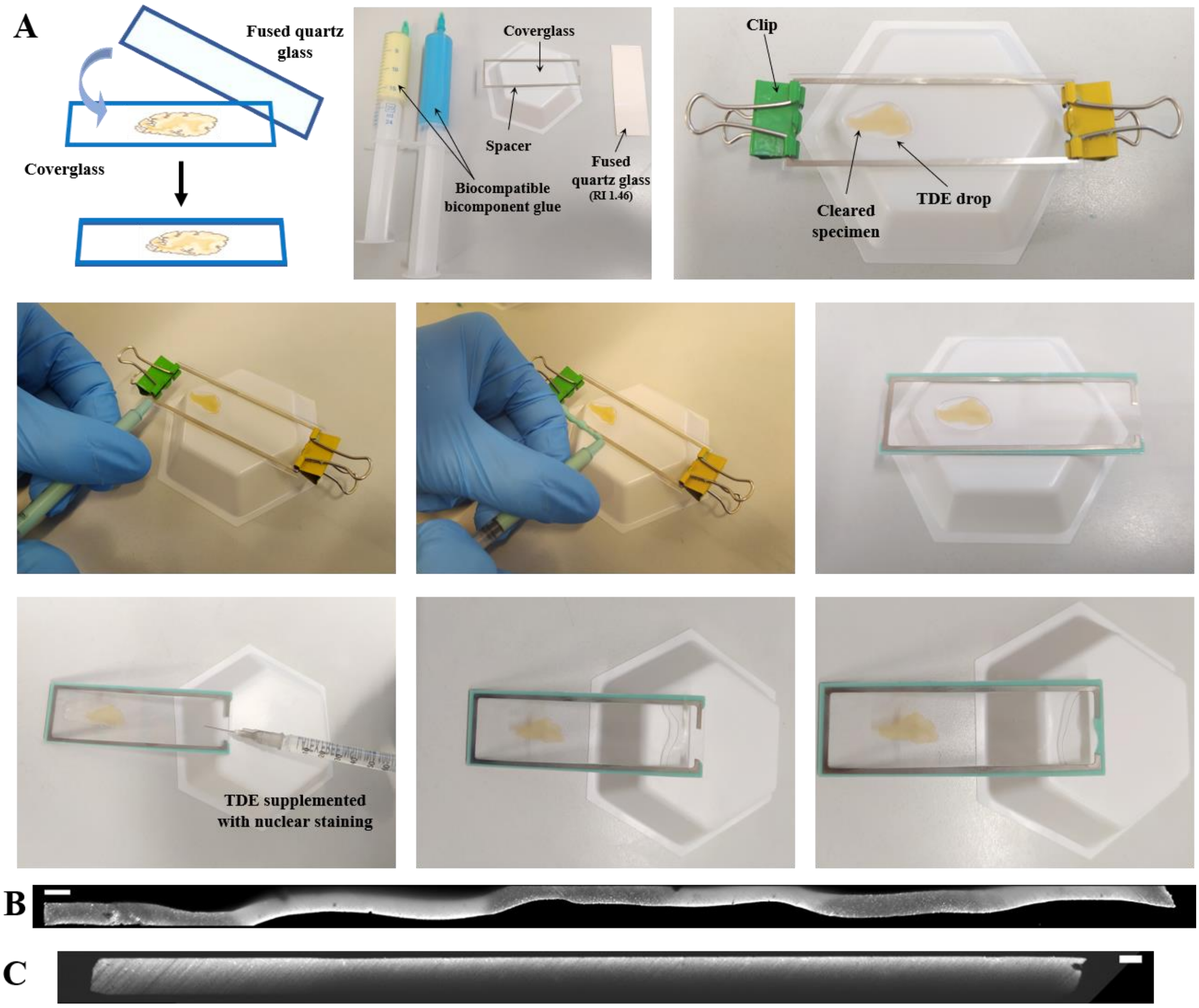


**Fig. S8. Sample holder assembly.** (A) The sandwich holder consists of a cover glass, a spacer (thickness of 500 um) and fused silica glass. The cleared specimen is fixed between the cover glass and the fused silica glass using a bicomponent glue. Such strategy allows soaking the sample in the TDE/PBS solution supplemented with a nuclear dye. Resliced images of mesoscopic reconstruction acquired by LSFM using the fused silica glass (B) or the sandwich apparatus (C). Scale bar:1 mm.

| Shrinking effect | PBS ($cm^2$) | SHORT ($cm^2$) |
|---|---|---|
| Sample 1 | 7.188 | 7.048 |
| Sample 2 | 7.181 | 6.92 |
| Sample 3 | 7.396 | 7.027 |

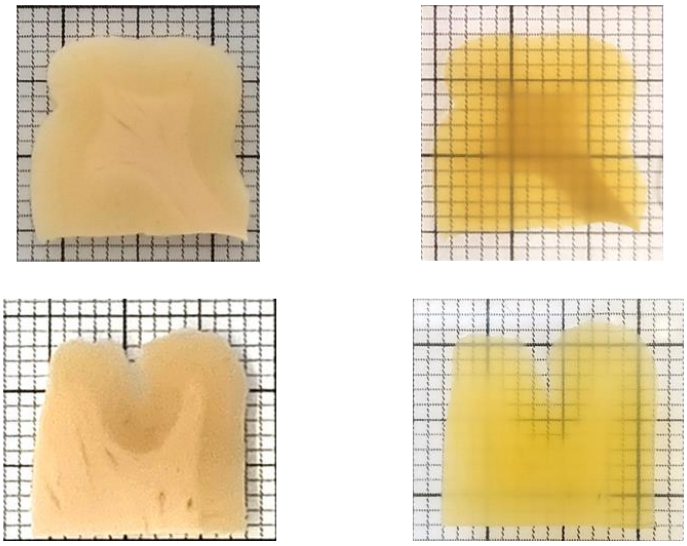

Shrinking characterization: (3.5 ± 1.5)%

**Fig. S9. Shrinking characterization of SHORT processed slices after incubation in the 68% TDE/PBS solution** (N = 3 samples). The slices to be enclosed into the holder, were equilibrated in the TDE solution, tuned for the refractive index matching of the delipidated sample (RI ≈ 1.46 with TDE/PBS solution at 68%), minimizing optical aberration and making the sample almost completely transparent. After equilibration of the processed sample into the PBS/TDE solution, we observed an isotropic tissue shrinkage of a factor of 3.5 ± 1.5 due to the RI matching obtained with the organic solvent TDE. Generally, the TDE solution is supplemented with nuclear staining.

| Molecule | Company | Cat. n. | Host | P/M | Dilution |
|---|---|---|---|---|---|
| NeuN | Merck | ABN91 | Chicken | P | 1:50 |
| GAD67 | Santa Cruz | Sc-28376 | Mouse | M | 1:200 |
| PV | Abcam | ab11427 | Rabbit | P | 1:200 |
| PV | Abcam | ab32895 | Goat | P | 1:200 |
| CB | Abcam | ab207528 | Rabbit | M | 1:200 |
| VIP | Abcam | ab214244 | Rabbit | M | 1:200 |
| SST | Abcam | ab30788 | Rat | M | 1:200 |
| NPY | Abcam | ab6173 | Sheep | P | 1:200 |
| NPY | Abcam | ab11247 | Mouse | M | 1:200 |
| SMI-32 | Merck | NE1023 | Mouse | M | 1:200 |
| SMI-31 | Eurogentec OptimAb | SMI-31P-050 | Mouse | M | 1:300 |
| Neurofilament | Abcam | ab4680 | Chicken | P | 1:200 |
| GluS | Merck | MAB302 | Mouse | M | 1:200 |
| MAP2 | Abcam | ab5392 | Chicken | P | 1:200 |
| GFAP | Abcam | ab194324 | Rabbit | M | 1:200 |
| Iba1 | Abcam | ab195031 | Rabbit | M | 1:200 |
| Coll IV | Abcam | ab6586 | Rabbit | P | 1:200 |
| Vim | Abcam | ab8069 | Mouse | M | 1:200 |
| CR | Proteintech | 66496-1-Ig | Mouse | M | 1:200 |
| CR | Proteintech | 12278-1-AP | Rabbit | P | 1:200 |
| Anti-Rat IgG, AF 568 | Abcam | ab175475 | Donkey | P | 1:200 |
| Anti-Rabbit IgG, AF 568 | Abcam | ab175470 | Donkey | P | 1:200 |
| Anti-Chicken IgY, AF 568 | Abcam | ab175711 | Goat | P | 1:200 |
| Anti-Mouse IgG, AF 568 | Abcam | ab175700 | Donkey | P | 1:200 |
| Anti-Sheep IgG, AF 568 | Abcam | ab175712 | Donkey | P | 1:200 |
| Anti-Rabbit IgG, AF 488 | Abcam | ab150077 | Goat | P | 1:200 |
| Anti-Chicken IgY, AF 488 | Abcam | ab150169 | Goat | P | 1:200 |
| Anti-Rabbit IgG, AF 488 | Jackson | 611-545-215 | AffiniPure Alpaca | P | 1:200 |
| Anti-Goat IgG, AF 488 | Jackson | 805-545-180 | AffiniPure Bovine | P | 1:200 |
| Anti Rabbit IgG, AF 647 | Jackson | 611-605-215 | AffiniPure Alpaca | P | 1:200 |
| Anti Mouse IgG, AF 647 | Abcam | ab150107 | Donkey | P | 1:200 |
| Anti Chicken IgY, AF 647 | Abcam | ab150171 | Goat | P | 1:200 |
| DAPI (Dilactate) | Thermo Fisher Scientific | D3571 | | | 1:50 |
| SYTOX GREEN | Thermo Fisher Scientific | S7020 | | | 1:100 |
| Propidium Iodide | Thermo Fisher Scientific | P3566 | | | 1:50 |

**Table S1. Antibodies and dyes compatible with SHORT and the used dilutions.**

| | Area | Age | Fixative | Fixation |
|---|---|---|---|---|
| **Sample 1** | - Precentral gyrus<br>- Hippocampus | 99 years old | 10% Formalin | 6 months |
| **Sample 2** | - Prefrontal cortex | 62 years old | 10% Formalin | 4 years |
| **Sample 3** | - Broca's area<br>- Motor cortex | 79 years old | 10% Formalin | 10 years |

**Table S2. Area, age, fixative molecule, and fixation of the human brain material used in this work.**
